# Structure-function studies on the serine protease domain of Factor VII activating protease (FSAP)

**DOI:** 10.1101/2025.11.25.690424

**Authors:** Sai Priya Sarma Kandanur, Bjørn Dalhus, Jonas Emsley, Sandip M. Kanse

**Affiliations:** Institute of Basic Medical Sciences, University of Oslo, Norway; Institute of Clinical Medicine, University of Oslo, Norway; Clinic for Laboratory Medicine, Oslo University Hospital, Norway; School of Pharmacy, University of Nottingham, Nottingham, UK

**Keywords:** HABP2, FSAP, serine protease, inhibitors, mutations, substrates

## Abstract

Factor VII activating protease (FSAP) is a circulating serine protease involved in stroke pathophysiology. This effect is likely mediated by the cleavage of substrates such as fibrinogen and extracellular histones. Our goal was to modify FSAP properties through mutagenesis to explore its different functions. Therefore, we expressed the serine protease domain (SPD) of FSAP with mutations and measured the cleavage of chromogenic substrate (S-2288), pro-urokinase (uPA), histones, and fibrinogen, as well as their inhibition by aprotinin, tissue factor pathway inhibitor (TFPI), and C1 inhibitor (C1Inh). Mutations are denoted using chymotrypsin numbering. Mutants KRHK63-65SSSS and K63aA/R63cA in the 60-loop were resistant to TFPI but not to aprotinin or C1Inh. Cleavage of pro-uPA and fibrinogen decreased, while histone and S-2288 cleavage remained normal. Mutants E96S, EDE96-99SSS, and D173A/E99S in the 99-loop produced a protease with reduced cleavage of histones and fibrinogen. EDE96-99SSS showed resistance to aprotinin, TFPI, and C1Inh, had lower affinity for S-2288, and exhibited less pro-uPA activation. The Q192D mutation was resistant to inhibition by aprotinin, TFPI, and C1Inh, but substrate cleavage remained unaffected. E218A was resistant to C1Inh inhibition but not to aprotinin or TFPI and showed normal substrate cleavage. R84S was resistant to aprotinin and TFPI but not C1Inh, with normal substrate cleavage. TFPI and C1Inh inhibited full-length FSAP more effectively than SPD-FSAP, though cleavage of histones and fibrinogen was more efficient with SPD-FSAP. These findings will enhance the understanding of the functional landscape of SPD-FSAP and help in developing mutants with altered properties to investigate FSAP functions.

## INTRODUCTION

Factor VII activating protease (FSAP) is a circulating serine protease(1) which belongs to the Clan PA and the family S1A of proteases, and emerged first approximately 600 million years ago in jawless vertebrates(2). It has high homology to urokinase-type plasminogen activator (uPA) and tissue plasminogen activator (tPA). Hepatocytes produce it(3) in its zymogen form and is encoded by the *hyaluronic acid binding protein 2 (HABP2)* gene(4). Pro-FSAP is activated to an enzymatically active two-chain FSAP comprising a heavy-chain and a light-chain (serine protease domain, SPD) that are linked by a disulfide bond. This activation is facilitated by extracellular histones released due to cell death or through the secretion of neutrophil extracellular traps (NETosis)(5). A naturally-occurring single nucleotide polymorphism in *HABP2* (Marburg I) (G534E) (G221E, chymotrypsin numbering) renders the protease completely inactive(6) and confers a higher risk for diseases such as carotid stenosis(7) and stroke(8). Studies on FSAP-knockout mice support the concept that FSAP is involved in the pathophysiology of thrombosis and stroke (9, 10), but not in normal development, growth, reproduction, or aging.

A preference for Arg-rich sequences was identified in FSAP substrates using short synthetic peptides and peptide phage display(11). Many protein substrates have been verified, including fibrinogen(12) and histones(5), but their *in vivo* significance is not fully elucidated. FSAP inhibitors include the α1-proteinase inhibitor, α2-anti-plasmin, anti-thrombin, TFPI, C1-Inhibitor (C1Inh), and plasminogen activator inhibitor-1 (PAI-1)(13–18).

Substrate and inhibitor specificity in serine proteases depends on the unique characteristics of their substrate-binding pockets as well as the eight surface loops located around the enzyme’s active site(19). These loops are called the activation, 37, 60, 75, 99, 148 (or autolysis loop), 170, 186 (or oxyanion stabilizing loop), and the 220 loop (or S1-entrance frame)(20). The heavy-chain also dictates substrate and inhibitor specificity as well as regulates enzyme activity in several ways. To date, no structural information or mutagenesis-based mapping data on FSAP activity is available. Using information from homology modeling and sequence alignment of related proteases, we introduced point mutations in various loops to examine the effects on enzymatic activity, substrate, and inhibitor selectivity.

We have recently demonstrated that the recombinant SPD-FSAP improves the outcome of ischemic stroke in mouse models(21) whereas other studies demonstrate complex actions in ischemic stroke(22, 23). The mutants can be used to study the biological role of FSAP, to engineer increased resistance to inhibitors, and to alter its specificity to enhance its stroke-protective effects.

## METHODS

### Homology modeling of WT-SPD-FSAP and sequence alignment with related proteases

The 247 residues of SPD-FSAP at the C-terminus (Ile 16- Phe 263, chymotrypsin numbering) was used as the input sequence to generate a homology model of active SPD-FSAP based on tPA (PDB ID- 5BRR: B), with which it has a 42% homology, using SWISS-MODEL (Fig. 1A). Positions of key features as well as the charge distribution is shown in Fig. 1B. The model had a QMEANDisCo global score of 0.68 ± 0.05 (range 0-1, with 1 being good) and GMQE Global modality quality estimation) score of 0.7 (range 0-1, with 1 being good), indicating good reliability of the prediction. To target specific residues in SPD-FSAP for mutations (Fig. 1C), we performed sequence alignment to identify non-conserved residues among related proteases. The residues selected for mutagenesis were then cross-validated with the AlphaFold model of full-length (FL) FSAP (Uniprot Q14520) (https://AlphaFoldAlphaFold.ebi.ac.uk/), in its zymogen form, to rule out residues involved in inter-domain interactions as well as exhibiting strong co-evolution (Table S1). Sequence alignment of SPD-FSAP and its related proteases: chymotrypsin, trypsin, thrombin, factor VII, factor X, factor XI, factor XII, activated protein C, kallikrein, plasminogen, urokinase-type plasminogen activator (uPA), and tissue-type plasminogen activator (tPA) was done using UniProt with default settings (Fig. S1). The charge distribution representations were made using PyMol Molecular Graphics System (version 2.4.0). Sequence alignment with 81 vertebrate species was performed and is presented as a conservation chart made using ConSurf; similarly, alignment with 100 different human paralogs was also performed (Fig. S2).

**Fig. 1:**
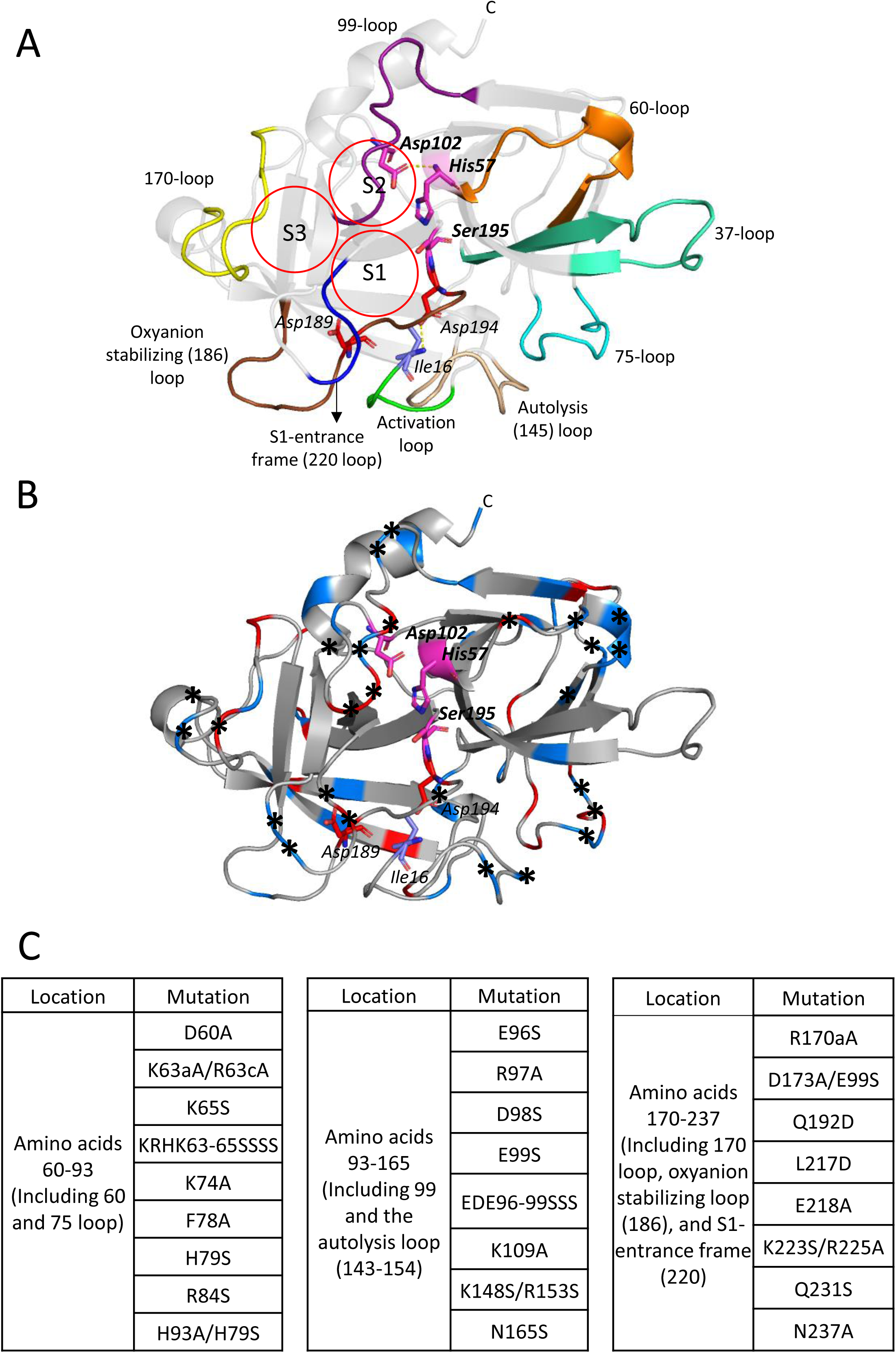
Predicted structure of SPD-FSAP and the position of the point mutations: **A.** Homology model of active SPD-FSAP (based on tPA, PDB ID 5BRR:B) showing the catalytic triad (magenta), the activation loop (green), the 37-loop (teal), 60-loop (orange), 705-loop (cyan), 99-loop (violet), the autolysis 148-loop (wheat), 170-loop (yellow), oxyanion stabilizing loop (brown), and the S1-entrance frame-200 loop (blue). The substrate specificity pockets S1, S2, and S3 (circled in red) and the salt bridge/ hydrogen bond (yellow dashed lines) between Ile 16-Asp194 and Asp102-His57, respectively, are indicated. **B.** The same model is shown in a ribbon form with the charged residues highlighted in red (negative) and blue (positive). Asterisk indicates the mutations introduced. **C.** An overview of all mutations introduced and their location in SPD-FSAP. The chymotrypsin numbering system is used.

### Expression, purification, and refolding of SPDs

PCR was used to introduce point mutations into the pASK-IBA33plus vector containing the gene coding for the SPD-FSAP (Fig. 1C), in addition to a C-terminal 6XHis-tag as described before(24). Protein expression and refolding were performed as described before(25) except that the 1 M NDSB-201 (3-(1-Pyridinio)-1-propanesulfonate) (1M) in the refolding buffer was replaced by arginine (1M). The inclusion body preparations were solubilized, refolded without purification, concentrated, buffer-exchanged into PBS, and stored at-80°C (crude SPDs). Thereafter, additional experiments were performed with selected constructs purified over Ni-NTA as described before(24). Protein expression was analyzed by SDS-PAGE under reducing conditions with Coomassie blue staining. The activation from the zymogen form to the active form is indicated by a shift in MW from 32 to 28 kDa under reducing conditions.

### Enzyme kinetics of SPDs and plasma FL-FSAP

FL-FSAP, isolated from human plasma (Dr. Michael Etscheid, Langen, Germany) and SPDs (300 nM) in Tris (25 mM, pH 7.4), NaCl (140 mM) (TBS) with 2 mM CaCl_2_ and 0.01% (v/v) Tween-20, were incubated with 0-0.3 mM chromogenic substrate, S-2288 (H-D-isoleucyl-L-prolyl-L-arginine-p-nitroanilinedihydro-chloride) (Chromogenix, Mölndal, Sweden). Maximal velocity for hydrolysis of S2288 was obtained by measuring the absorbance at 405 nm every min for a period of 1 hour at 37°C(24). The data were fitted into the Michaelis-Menten equation using GraphPad Prism Software, version 8.3.0 from GraphPad software (San Diego, CA).

### Activation of pro-uPA

Increasing concentrations (0-416 nM) of pro-urokinase (pro-uPA) were incubated with 0.03 µg/ml of FL-FSAP and SPDs for 20 min at 37°C, and the activation of pro-uPA was measured by the addition of 0.2 mM chromogenic substrate S-2444 (L-pyroglutamyl-glycyl-L-arginine-p-nitroanilinedihydro-chloride) (Chromogenix). The absorbance was measured every min at 405 nm for 60 min at 37°C in a plate reader.

### Active site titration with aprotinin and inhibition assays with aprotinin, C1Inh, and TFPI

The percentage active fraction was estimated as previously described(26). SPDs were incubated with increasing amounts of aprotinin (Sigma, Oslo, Norway) (0-1000 nM), and chromogenic substrate turnover was measured. An initial linear decrease in velocity with increasing concentration of aprotinin was plotted against aprotinin:SPD molar ratios. Mutants that exhibited altered inhibition with aprotinin cannot be titrated using this method, in which case an average value based on the other mutants was used. Thus, these results need to be reconfirmed with other methods.

FL-FSAP and SPDs (300 nM) were incubated with increasing concentrations (0-1000 nM) of C1Inh (Pharming Group, Leiden, Netherlands), Aprotinin, and TFPI (Prof. Tilman Hackeng, Maastricht, Netherlands) for 30 min at room temperature, and then S2288 (0.5 mM) turnover was measured as described above. IC_50_ values were obtained using GraphPad, and percentage residual activity at 1000 nM of inhibitor was noted.

### Cleavage of histones and fibrinogen

Histone mix (Calf thymus, Sigma, Oslo, Norway) and plasminogen-depleted fibrinogen (Merck Millipore, Darmstadt, Germany) (2 mg/ml) were incubated with 0-833 nM of SPDs in TBS with 2 mM CaCl_2_. We used a natural mixture of calf thymus histones so that their normal complexation state is maintained, and we could capture all types of cleavage simultaneously (27). Each reaction was incubated at 37°C for 60 min, and the reaction was stopped using non-reducing SDS sample loading buffer. Similarly, 2 mg/ml of histones or fibrinogen was incubated with 833 nM FL-FSAP or SPDs for 1, 5, 10, 20, 30, 60, and 100 minutes, respectively. The samples were run on 4-20 % gradient SDS-PAGE gels, followed by Coomassie Blue staining, and densitometric analyses of the bands using Image J software were performed. IC_50_ and Time_50_ values were obtained using GraphPad, and percentage cleavage at the highest concentration of SPDs was noted.

### Replication and statistics

Results are representative of 3 independent experiments and are expressed as mean ± SD (n= independent experiments). Histone and fibrinogen degradation assays were performed in two independent experiments and are presented as mean ± range. Statistical analysis was performed using one-way ANOVA followed by Dunnett’s test using Graphpad.

## RESULTS

### Expression of SPDs and their active site titration

Most mutant SPDs underwent zymogen activation during the refolding process, resulting in a 28 kDa protein band (Fig. 2A), while mutants D60A, KRHK63-65SSSS, K65S, H93A/H79S, E96S, K109A, D173A/E99S, K148S/R153S, and Q192D were partially in the zymogen form (32 kDa) (Fig. S3). MI-mutant (G221E) was completely in the zymogen form, as were the mutants L156K/Q192D, G193A/W, L217W/E218W, and G216A (data not shown). Some mutants, E76A and K186A, were not expressed at all. Active-site titration of crude SPDs with aprotinin resulted in WT-SPD with an active fraction of 2.4%, EDE96-99SSS, K63aA/R63cA, R170A, and L217D had an active fraction ranging between 7-7.5% and K223S/R225A was 15% active, whilst the others were in the range 1-4%. In comparison, human plasma-purified FL-FSAP had 80% of active protease. For mutants not inhibited by aprotinin (see later), we estimated an average refolding based on the other mutants. In an earlier study, using an NDSB-201-based refolding buffer, we observed that 50% of SPD was correctly refolded, but there was strong precipitation. In the current study, we used arginine in the refolding buffer, which resulted in no precipitation, but this also decreased the overall percentage of active refolded protein. Further purification of SPDs over Ni-NTA and/ or benzamidine Sepharose increased purity but also increased autoproteolysis.

**Fig. 2:**
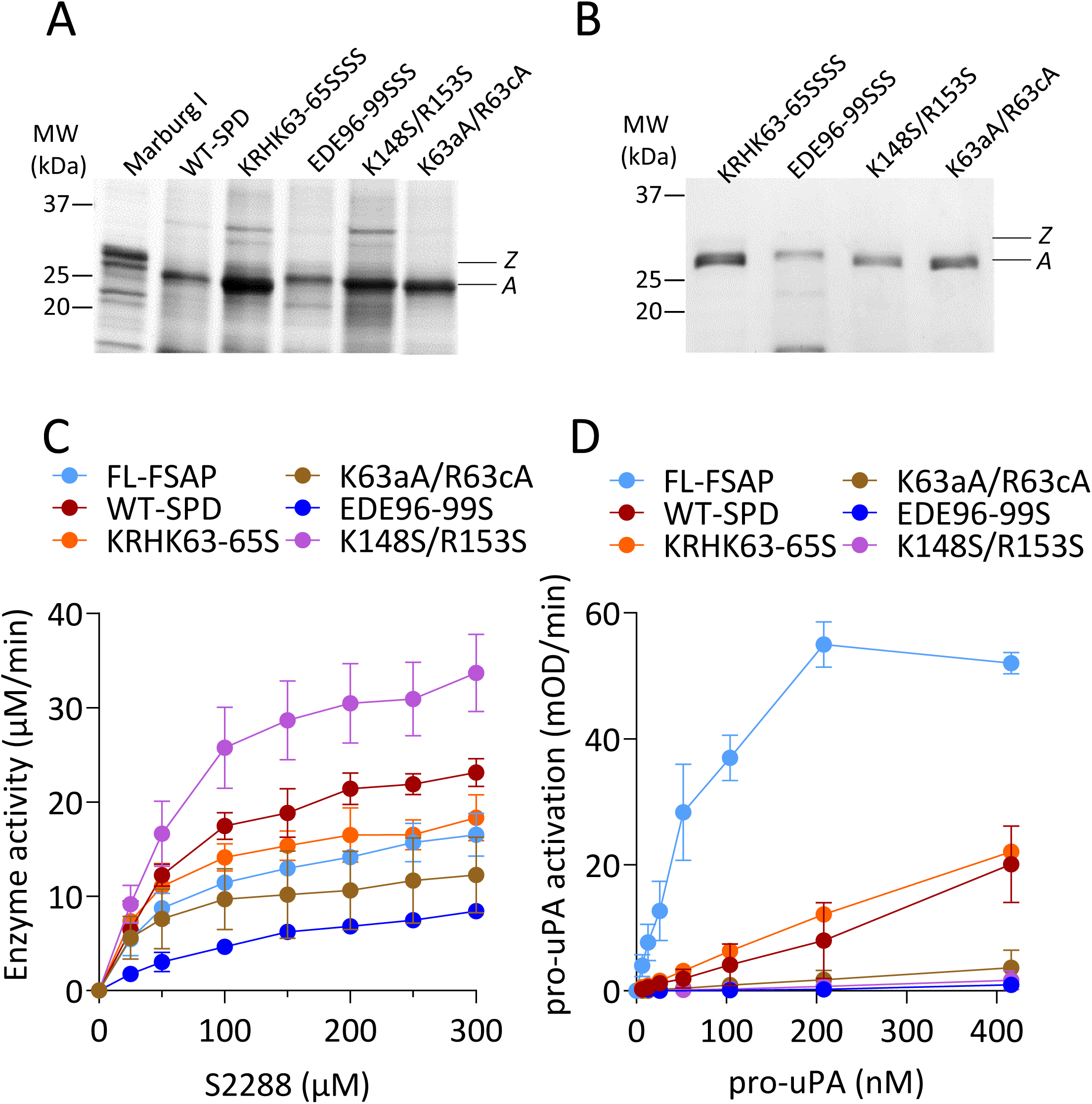
SDS-PAGE analysis of SPD mutants and their activity against chromogenic substrate, S2288, and physiological substrate, pro-uPA: **A.** SDS-PAGE gel showing zymogen form (*Z*, 32 kDa) and the active form (*A,* 28 kDa) of SPDs for selected mutants in crude refolded extracts. **B.** SDS-PAGE gel of SPDs purified over a Ni-NTA column. **C.** Michaelis-Menten plot of FL-FSAP, WT-SPD, and selected mutants showing the cleavage of S2288 by the SPDs (300 nM). Substrate hydrolysis was measured as absorbance at 405 nm and is represented as μM/min. The Michaelis-Menten constants for all mutants are in Table 1. **D.** pro-uPA (0-420 nM) was activated by 0.03 µg/ml FL-FSAP and purified SPDs for 20 min at 37°C. The activity of uPA was measured by the hydrolysis of chromogenic substrate S-2444 and is represented as mOD/min. The value of the slope of mOD/min/ nM pro-uPA is shown in Table 2. All values are mean ± SD (n=3).

**Table 1:** Michaelis-Menten constants for plasma FL-FSAP and SPDs using chromogenic substrate S2288. Values higher (blue) and lower (red) than WT are indicated. All values are mean ± SD (n=3),* indicates p < 0.05.

| FL-FSAP or SPD | $K_{cat}$<br>(min<sup>-1</sup>) | $K_m$<br>(μM) | $K_{cat}/K_m$ (min<sup>-1</sup><br>μm<sup>-1</sup>) |
| --- | --- | --- | --- |
| FL-FSAP | 952 ± 224 | 78 ± 45 | 15 ± 9 |
| WT-SPD | 1347 ± 126 | 76 ± 30 | 19 ± 6 |
| D60A | 776 ± 115* | 38 ± 11 | 21 ± 5 |
| K63aA/R63cA | 620 ± 231* | 41 ± 17 | 18 ± 11 |
| K65S | 2491 ± 473* | 94 ± 46 | 31 ± 14 |
| KRHK63-65SSSS | 940 ± 132 | 44 ± 14 | 22 ± 5 |
| K74A | 1215 ± 207 | 60 ± 45 | 26 ± 11 |
| F78A | 2392 ± 149* | 109 ± 24 | 22 ± 3 |
| H79S | 768 ± 221* | 44 ± 15 | 18 ± 2 |
| R84S | 1077 ± 128 | 58 ± 21 | 20 ± 7 |
| H93A/H79S | 1894 ± 489* | 105 ± 30 | 18 ± 2 |
| E96S | 346 ± 69* | 46 ± 28 | 9 ± 4 |
| R97A | 636 ± 184* | 43 ± 17 | 15 ± 2 |
| D98S | 679 ± 163* | 55 ± 15 | 13 ± 2 |
| E99S | 891 ± 194 | 45 ± 12 | 20 ± 1 |
| EDE96-99SSS | 612 ± 106* | 183 ± 94* | 4 ± 1* |
| K109A | 934 ± 104 | 47 ± 15 | 21 ± 5 |
| K148S/R153S | 1948 ± 171 | 76 ± 14 | 27 ± 7 |
| N165S | 1030 ± 180 | 48 ± 18 | 23 ± 5 |
| R170aA | 645 ± 84* | 48 ± 6 | 14 ± 3 |
| D173A/E99S | 110 ± 3* | 35 ± 9 | 3 ± 1* |
| Q192D | 1335 ± 58 | 52 ± 6 | 26 ± 4 |
| L217D | 883 ± 20 | 93 ± 24 | 10 ± 2 |
| E218A | 1646 ± 85 | 115 ± 31 | 15 ± 3 |
| K223S/R225A | 506 ± 44* | 19 ± 4 | 27 ± 5 |
| Q231S | 1382 ± 33 | 50 ± 4 | 28 ± 3 |
| N237A | 1409 ± 203 | 61 ± 14 | 24 ± 6 |

**Table 2:** Pro-uPA activation by plasma FL-FSAP and SPDs. The values chromogenic substrate turnover/ pro-uPA concentration were used for comparing the mutants. All values are mean ± SD (n=3), * indicates p < 0.05.

| FL-FSAP or SPD | Slope ± SD |
| --- | --- |
| FL-FSAP | $3 \pm 0.2$ |
| WT-SPD | $1 \pm 0.3$ |
| KRHK63-65SSSS | $1 \pm 0.05$ |
| EDE96-99SSS | $0.05 \pm 0.03^*$ |
| K148S/R153S | $0.09 \pm 0.07^*$ |
| K63aA/R63cA | $0.2 \pm 0.1^*$ |

### Proteolytic activity against chromogenic substrate S-2288

Michaelis-Menten analysis with a small peptidic chromogenic substrate, S-2288, showed that FL-FSAP and WT-SPD had similar *K_m_* and *K_cat_* values as reported earlier(25). Representative data on the mutants are in Fig. 2C, and the constants are in Table 1. Overall, there were small changes in the enzymatic properties of these mutants. *K_m_* was significantly increased for the EDE96-99SSS mutant only. Catalytic efficiency was decreased only for the DE96-99S and D173A/E99S mutants. K*_cat_* was moderately altered for many mutants (Table 1). Thus, WT-SPD and FL-FSAP had comparable kinetic properties, and the mutations had weak effects on affinity, except for the EDE96-99SSS mutant.

### Activation of pro-uPA

We planned to do these experiments with the crude preparations from all the constructs, but sufficient protein was not available for these studies. We compared the activation of pro-uPA(16) with Ni-NTA-purified WT-SPD, KRHK63-65SSSS, K63aA/R63cA, EDE96-99SSS, and K148S/R153S (Fig. 2B). With increasing concentrations of pro-uPA, the cleavage with FL-FSAP reached saturation; however, this was not the case with the SPDs (Fig. 2D). For quantitative comparison, we calculated the slopes of the velocity/ pro-uPA concentration curves (Fig. 2D, Table 2). WT-SPD was a weaker activator of pro-uPA compared to FL-FSAP, and the mutants EDE96-99SSS, KRHK63-65SSSS, and K148S/R153S were weak activators compared to WT-SPD.

### Inhibition with Aprotinin, TFPI, and C1Inh

The inhibition of the mutants by different types of inhibitors was examined using S2288 chromogenic substrate assays. Kunitz-type inhibitors like aprotinin and TFPI inhibit FSAP(28). The inhibition of FL-FSAP and WT-SPD with aprotinin (Fig. 3A) gave similar IC_50_ values (3 and 5 nM, respectively), but with TFPI, the IC_50_ values were 28 ± 6 nM and 220 ± 146 nM, respectively (Table 3). This confirms the earlier report that the heavy-chain of FSAP plays an important role in its interaction with TFPI(28). Mutants R84S, EDE96-99SSS, and Q192D exhibited resistance to inhibition by aprotinin and TFPI. Mutants K63aA/R63cA and KRHK63-65S were resistant to TFPI, but not aprotinin.

**Fig. 3:**
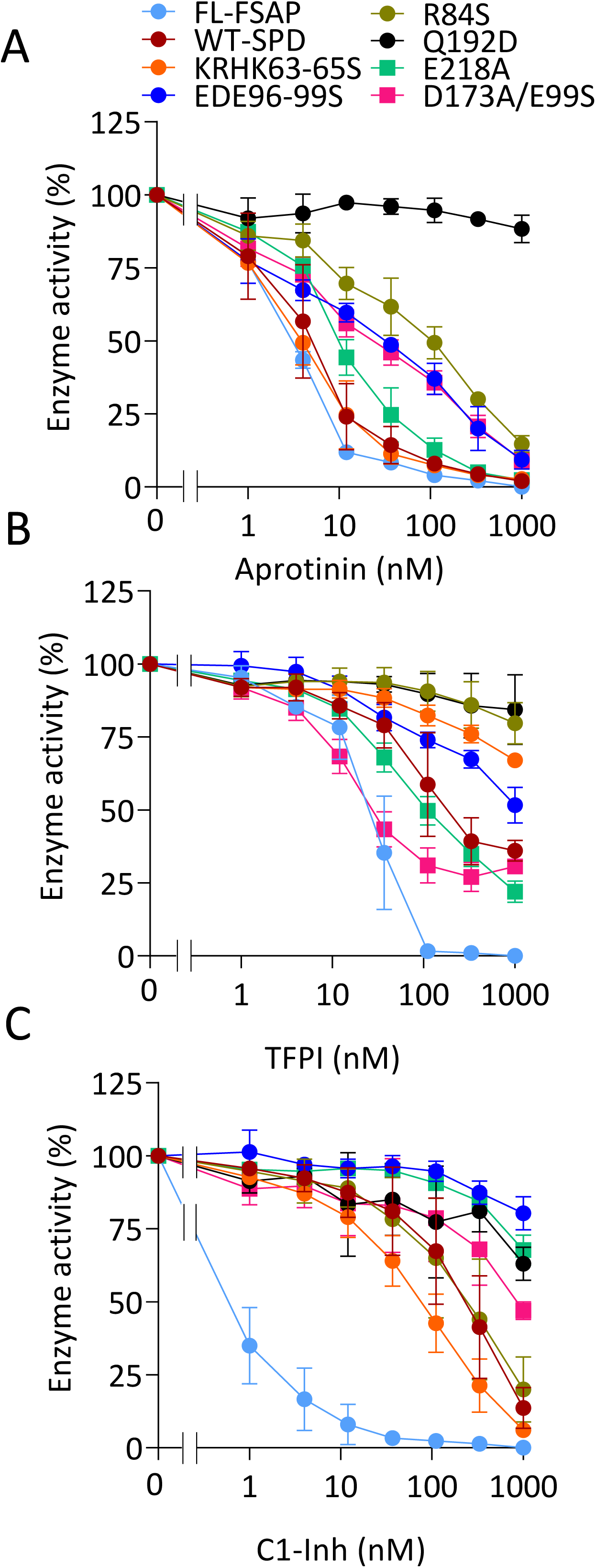
Inhibition of SPD-FSAP mutants by Aprotinin, TFPI, and C1Inh: FL-FSAP and SPD mutants (300 nM) were incubated at RT for 30 minutes with 0-1000 nM of aprotinin **(A)**, TFPI **(B)**, and C1Inh **(C)**. Inhibition of FSAP and SPDs was measured by the hydrolysis of chromogenic substrate S2288 and is represented as % enzyme activity. All values are mean ± SD (n=3). This figure is a representation of the experiment carried out on all mutants. The IC_50_ values for all mutants are shown in Table 3.

**Table 3:** IC_50_ values of plasma FL-FSAP and SPDs against Aprotinin, C1Inh, and TFPI using chromogenic substrate S2288. Values higher (blue) and lower (red) than WT are indicated. All values are mean ± SD (n=3), * indicates p < 0.05.

| Mutant | IC<sub>50</sub> (nM) |  |  | % Residual activity at 1000 nM |  |  |
| --- | --- | --- | --- | --- | --- | --- |
|  | Aprotinin | TFPI | C1Inh | Aprotinin | TFPI | C1Inh |
| FL-FSAP | 3 ± 0.2 | 28 ± 6 | 11 ± 4 | 0 | 0 | 0 |
| WT-SPD | 5 ± 3 | 220 ± 146 | 423 ± 191 | 2 ± 1 | 36 ± 4 | 14 ± 7 |
| D60A | 23 ± 18 | 82 ± 35 | 354 ± 13 | 7 ± 3* | 31 ± 4 | 12 ± 8 |
| K63aA/R63cA | 4 ± 6 | >1000* | 255 ± 114 | 2 ± 0 | 57 ± 7* | 5 ± 3 |
| K65S | 5 ± 1 | 688 ± 280* | 592 ± 47 | 1 ± 1 | 43 ± 5 | 16 ± 3 |
| KRHK63-65SSSS | 4 ± 1 | >1000* | 210 ± 78 | 3 ± 1 | 67 ± 2* | 6 ± 1 |
| K74A | 9 ± 3 | 459 ± 49 | 187 ± 88 | 2 ± 1 | 44 ± 5 | 6 ± 2 |
| F78A | 14 ± 1 | 853 ± 133 | 292 ± 91 | 2 ± 1 | 46 ± 1 | 11 ± 4 |
| H79S | 7 ± 4 | 468 ± 420 | 406 ± 195 | 2 ± 1 | 38 ± 10 | 14 ± 7 |
| R84S | 95 ± 36* | >1000* | 446 ± 259 | 15 ± 3* | 80 ± 7* | 20 ± 11 |
| H93A/H79S | 9 ± 3 | 134 ± 119 | 450 ± 139 | 3 ± 1 | 26 ± 6 | 17 ± 6 |
| E96S | 36 ± 12 | 227 ± 163 | 645 ± 140 | 8 ± 0* | 39 ± 4 | 29 ± 6 |
| R97A | 12 ± 11 | 338 ± 257 | 410 ± 86 | 1 ± 1 | 30 ± 5 | 12 ± 4 |
| D98S | 15 ± 13 | 522 ± 525 | 620 ± 91 | 2 ± 1 | 45 ± 8 | 26 ± 6 |
| E99S | 15 ± 4 | 599 ± 216 | 254 ± 30 | 5 ± 1 | 47 ± 2 | 7 ± 3 |
| EDE96-99SSS | 31 ± 4 | >1000* | >1000* | 9 ± 3* | 52 ± 6* | 80 ± 6* |
| K109A | 16 ± 10 | 252 ± 248 | 373 ± 142 | 2 ± 1 | 33 ± 5 | 11 ± 7 |
| K148S/R153S | 11 ± 9 | 507 ± 347 | 465 ± 140 | 1 ± 1 | 43 ± 5 | 16 ± 3 |
| N165S | 7 ± 8 | 143 ± 84 | 300 ± 154 | 1 ± 1 | 32 ± 5 | 7 ± 6 |
| R170aA | 28 ± 23 | 215 ± 145 | 479 ± 164 | 3 ± 1 | 34 ± 5 | 17 ± 4 |
| D173A/E99S | 30 ± 15 | 29 ± 10 | >1000* | 9 ± 0* | 31 ± 3 | 47 ± 3* |
| Q192D | >1000* | >1000* | >1000* | 88 ± 5* | 84 ± 12* | 49 ± 24* |
| L217D | 24 ± 21 | 165 ± 36 | 445 ± 85 | 1 ± 1 | 27 ± 6 | 7 ± 2 |
| E218A | 12 ± 3 | 122 ± 34 | >1000* | 2 ± 1 | 22 ± 4 | 68 ± 5* |
| K223S/R225A | 49 ± 23* | 107 ± 30 | 218 ± 63 | 5 ± 2 | 25 ± 1 | 2 ± 1 |
| Q231S | 16 ± 7 | 300 ± 158 | 310 ± 95 | 3 ± 1 | 42 ± 8 | 8 ± 7 |
| N237A | 17 ± 9 | 180 ± 164 | 424 ± 80 | 2 ± 1 | 32 ± 7 | 17 ± 5 |

In addition to kunitz-type inhibitors, FSAP is also inhibited by serine protease inhibitors like C1Inh(13). Inhibition of FL-FSAP and WT-SPD with increasing concentration of C1Inh (Fig. 3C) gave an IC_50_ of 11 ± 4 nM and 423 ± 191 nM, respectively (Table 3), confirming the importance of the heavy-chain for this interaction. EDE96-99SSS, D173A/E99S, Q192D, and E218A exhibited reduced inhibition by C1Inh. Since both types of inhibitors target the active site, some mutations likely influence the inhibition by both types of inhibitors. However, E218A was an exception to this and was exclusively inhibited by C1Inh.

### Cleavage of histones and fibrinogen

The effect of mutants on the cleavage of large protein substrates was then tested. FSAP cleaves histones, thereby protecting cells from histone cytotoxicity(5). The cleavage of H1 and H2A/ H2B was more effective with WT-SPD than FL-FSAP (Table 4, Fig. 4A). Mutants E96S, EDE96-99SSS, and D173A/E99S exhibited reduced efficiency in H1 and H2A/H2B cleavage in terms of IC_50_ as well as the time taken to degrade 50% of substrate (T_50_). Although there was a difference in the IC_50_ values for H1 and H2A/H2B, the pattern of how the mutants cleaved both these proteins was identical. Due to their migration characteristics on SDS-PAGE, we could not quantify H3/H4 cleavage with this method.

**Fig. 4:**
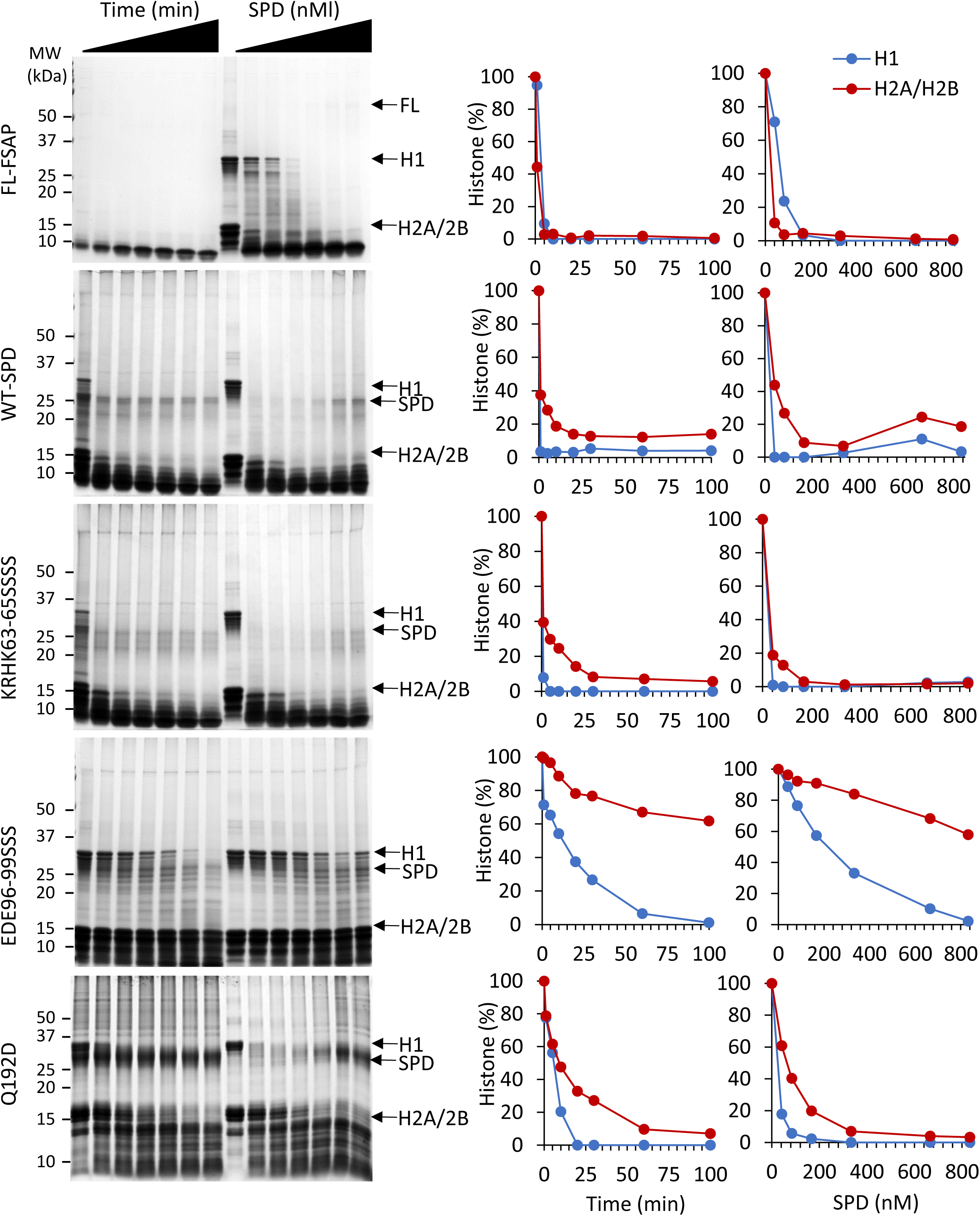
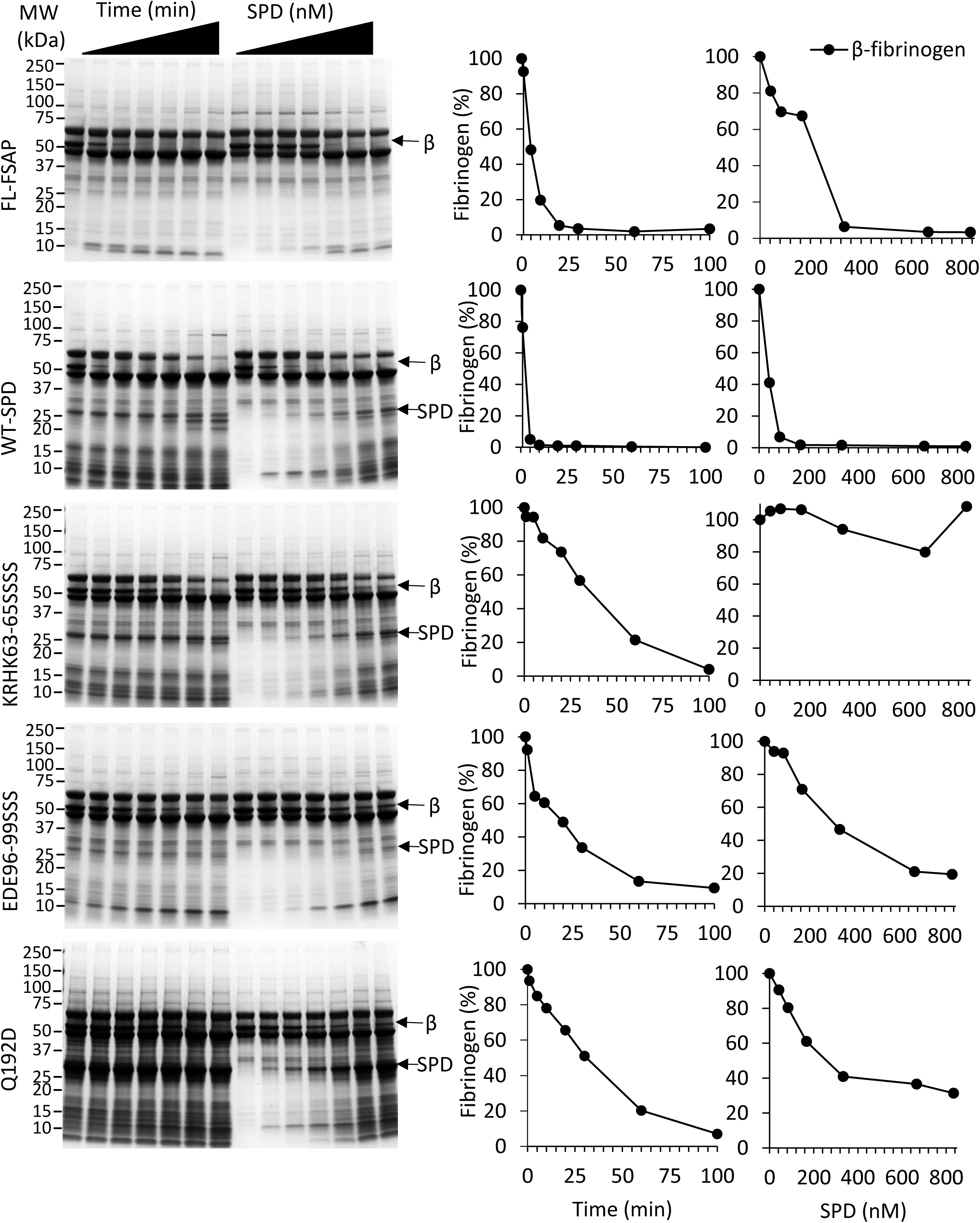
Effect of SPD mutants on histone and fibrinogen cleavage: **A.** SPD mutants (833 nM) were incubated with histones (2 mg/ml) for 1, 5, 10, 20, 30, 60, and 100 minutes at 37 °C. Increasing concentrations of SPD mutants (0-833 nM) were incubated with histones (2 mg/ml) at 37°C for 1 h and analyzed by SDS-PAGE. Migration of Histone H1, H2A/H2B, and SPDs is indicated. Densiometric analyses of residual histones against time and residual H1 and H2A/H2B are shown in the panels to the right. **B.** The assays for fibrinogen were carried out as described for histones. The migration of beta (β)-chain fibrinogen and SPD is indicated. Densiometric analyses of residual β-fibrinogen against time and residual β-fibrinogen are shown in the panel to the right. All values are mean ± range (n=2). This figure is a representation of the experiment carried out on all mutants in two independent experiments. Quantitative analysis of the data with all mutants is in Table 4.

**Fig. 5:**
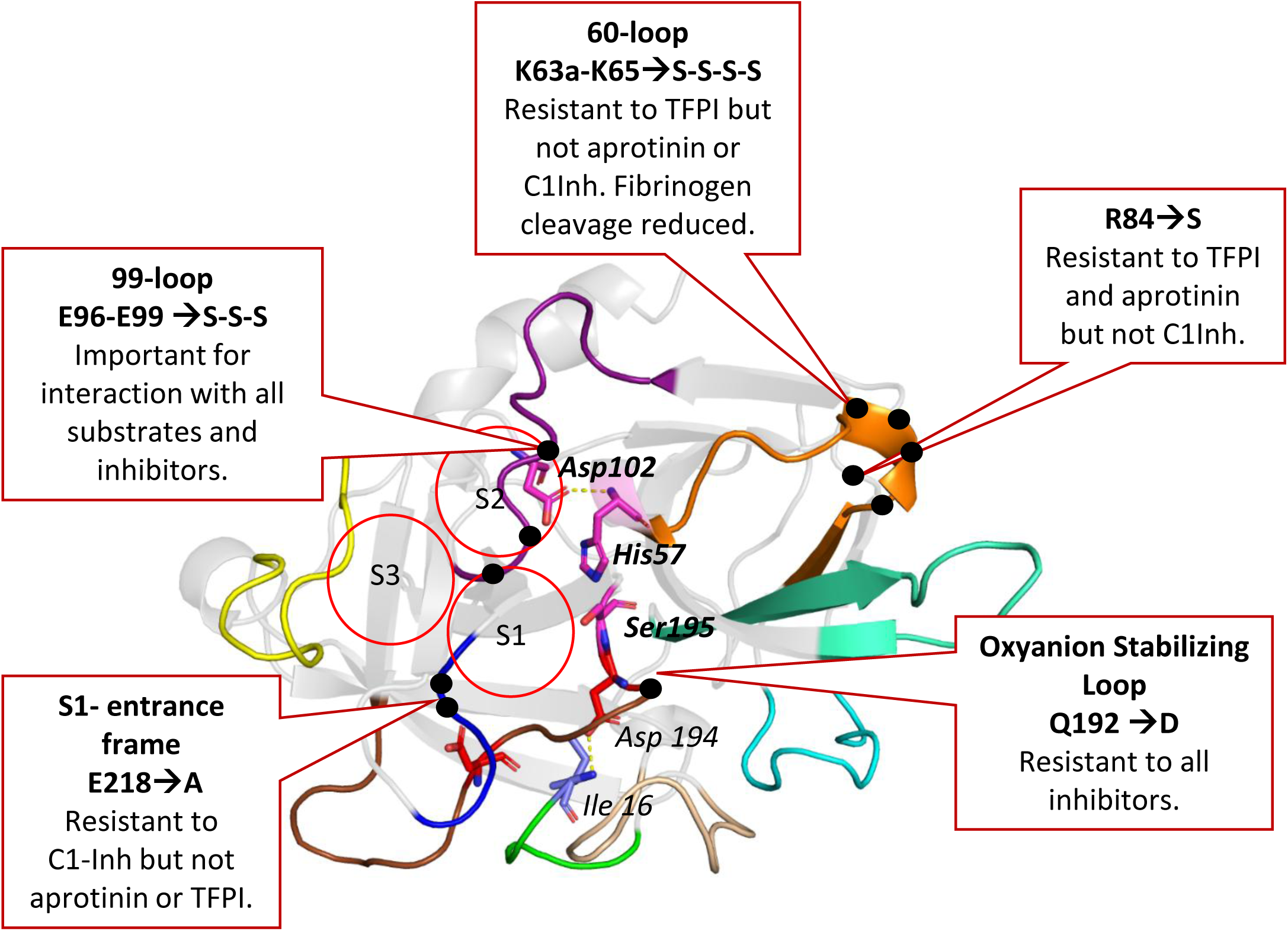
A homology model of SPD-FSAP summarizing the effects of key mutations on its interactions with substrates and inhibitors.

**Table 4:** Degradation of substrates by plasma FL-FSAP and SPDs. IC50 values and the time at which 50 % of the substrate is degraded are indicated. Values higher (blue) and lower (red) than WT are indicated. All values are mean ± range (n=2), * indicates p < 0.05.

| Mutant | IC<sub>50</sub> (μg/ml) | | | Degradation of 50% substrate (min) | | |
| --- | --- | --- | --- | --- | --- | --- |
|  | Histones |  | Fibrinogen | Histones |  | Fibrinogen |
|  | H1 | H2A/H2B |  | H1 | H2A/H2B |  |
| FL-FSAP | 2 ± 0.7 | 0.1 ± 0.1 | 6 ± 0.3 | 1 ± 2 | 0.5 ± 0.6 | 5 ± 0.4 |
| WT-SPD | 0.1 | 0.6 ± 0.4 | 0.1 | 1 ± 0.6 | 0.5 ± 0.2 | 1 ± 0.6 |
| D60A | 0.9 ± 0.02 | 1 ± 0.5 | 0.9 ± 0.02 | 1 ± 0.5 | 3 ± 1 | 1 ± 0.5 |
| K63aA/R63cA | 0.7 ± 0.08 | 2 ± 1 | >20* | 1 ± 0.2 | 2 ± 0.3 | 1 ± 0.2 |
| K65S | 0.4 ± 0.4 | 0.5 ± 0.2 | 0.4 ± 0.5 | 1 ± 0.1 | 1 ± 0.2 | 1 ± 0.1 |
| KRHK63-65SSSS | 0.3 ± 0.3 | 1 ± 1 | >20* | 1 ± 0.1 | 0.4 ± 0.2 | 1 ± 0.1 |
| K74A | 0.4 ± 0.4 | 1 ± 0.3 | 0.4 ± 0.5 | 1 ± 0.2 | 1 ± 0.2 | 1 ± 0.2 |
| F78A | 0.1 | 0.6 ± 0.1 | 0.1 | 1 ± 0.1 | 1 ± 0.2 | 1 ± 0.1 |
| H79S | 0.9 ± 0.02 | 1.0 ± 0.2 | 0.9 ± 0.02 | 2 ± 0.8 | 2 ± 1 | 2 ± 0.8 |
| R84S | 0.8 ± 0.08 | 0.7 ± 0.2 | 0.8 ± 0.08 | 1 ± 0.1 | 1 ± 0.1 | 1 ± 0.1 |
| H93A/H79S | 0.1 | 0.2 ± 0.2 | 0.1 | 1 ± 0.2 | 1 ± 0.4 | 1 ± 0.2 |
| E96S | 3.1 ± 0.03* | 6 ± 0.8* | 3 ± 0.03* | 9 ± 0.05* | 13 ± 1* | 9* |
| R97A | 0.1 | 0.5 ± 0.5 | 0.1 | 1 ± 0.1 | 1 ± 0.1 | 1 ± 0.1 |
| D98S | 0.4 ± 0.4 | 0.8 ± 0.04 | 0.4 ± 0.4 | 1 ± 0.3 | 1 ± 0.01 | 1 ± 0.3 |
| E99S | 0.4 ± 0.4 | 1 ± 0.2 | 0.4 ± 0.4 | 3 ± 4 | 3 ± 2 | 3 ± 4 |
| EDE96-99SSS | 9 ± 4* | >20* | 9 ± 4* | >100* | >100* | 18 ± 10* |
| K109A | 0.7 ± 0.2 | 0.8 ± 0.4 | 0.7 ± 0.2 | 1 ± 0.1 | 3 ± 3 | 1 ± 0.1 |
| K148S/R153S | 0.4 ± 0.4 | 0.6 ± 0.4 | 0.4 ± 0.4 | 1 ± 0.2 | 1 ± 0.1 | 1 ± 0.2 |
| N165S | 0.4 ± .04 | 0.8 | 0.4 ± 0.4 | 1 ± 0.1 | 1 ± 0.2 | 1 ± 0.1 |
| R170aA | 0.7 ± 0.3 | 2 ± 0.4 | 0.7 ± 0.03 | 2 ± 1 | 3 ± 0.5 | 2 ± 1 |
| D173A/E99S | >20* | >20* | >20* | >100* | >100* | >100* |
| Q192D | 0.4 ± 0.2 | 2 ± 0.1 | 0.4 ± 0.2 | 4 ± 3 | 8 ± 8 | 4 ± 3 |
| L217D | 0.7 ± 0.02 | 2 ± 1 | 0.7 ± 0.02 | 2 ± 0.5 | 4 ± 3 | 2 ± 0.5 |
| E218A | 0.7 ± 0.06 | 1 ± 0.6 | 0.7 ± 0.06 | 1 ± 0.6 | 5 ± 6 | 1 ± 0.6 |
| K223S/R225A | 0.9 ± 0.05 | 0.8 ± 0.03 | 0.9 ± 0.05 | 2 ± 0.6 | 3 ± 2 | 2 ± 0.6 |
| Q231S | 0.5 ± 0.5 | 0.5 ± 0.4 | 0.5 ± 0.5 | 1 ± 0.1 | 2 ± 0.5 | 1 ± 0.1 |
| N237A | 0.4 ± 0.4 | 0.6 ± 0.1 | 0.4 ± 0.4 | 1 ± 0.3 | 1 ± 0.4 | 1 ± 0.3 |

FSAP cleaves fibrinogen, particularly the β-chain but not the α- and ɣ-chain, and promotes the lysis of clots(12). FL-FSAP cleaved β-fibrinogen with lower efficacy than WT-SPD, suggesting that the heavy-chain of FSAP might have an inhibitory effect on this interaction (Table 4, Fig. 4B). The three mutants that exhibited reduced histone cleavage, E96S, EDE96-99SSS, and D173A/E99S, were also unable to cleave fibrinogen effectively. Additionally, mutants K63aA/R63cA and KRHK63-65SSSS exhibited diminished cleavage of fibrinogen. Thus, some mutants exhibited reduced cleavage of fibrinogen and histones, whereas the 60-loop mutants reduced cleavage of fibrinogen only, but none showed exclusive defects in histone cleavage only.

## DISCUSSION

The flexible surface loops in serine proteases play an important role in regulating substrate specificity, inhibition, zymogen activation, and interaction with cofactors(20). tPA and activated protein C (APC) are two serine proteases that have been altered to enhance their selectivity and efficacy as drugs to treat stroke. A substitution of Lys-His-Arg-Arg in the 37 loop of tPA increases its resistance to PAI-1 and its half-life i*n vivo*(29). Similarly, mutating Lys-Lys-Lys at position 191-193 just outside the SPD reduces the anti-coagulant effects of APC by 90%(30). In this report, we have introduced point mutations in SPD-FSAP to identify specific sites important for interactions with substrates and inhibitors and to dissociate the multiple roles of FSAP from one another. Mutations may also impact the allosteric interactions between the different loops that, in turn, will influence the function of FSAP. Furthermore, mutants may exhibit improved expression and stability, which will be important in elucidating the structure of FSAP.

### Mutations in the 60 Loop

Mutations in the 60-loop, K63aA/R63cA, and KRHK63-65SSSS exhibited reduced activation of pro-uPA, but cleavage of S2288 was normal. Both these mutants exhibited resistance to inhibition by TFPI. The 60-loop regulates specificity at the S2 pocket, and it mediates molecular recognition in APC(31). In thrombin, it acts like a lid that prevents the binding of less specific substrates(32). In tPA, the presence of an additional four residues in the 60-loop (amino acids 60a-c) has been shown to enhance specificity for plasminogen and PAI-1, while limiting binding to other substrates(33). Similarly, there are four additional residues in the 60-loop of FSAP, three of which are positively charged. Hence, it is highly likely that this region is important for FSAP’s selectivity. This mutant can be used to test the effects of FSAP on fibrinolysis as well as the role of its interaction with TFPI.

### Mutations in the 99 Loop

The mutant EDE96-99SSS exhibited reduced cleavage of S2288, histones, and fibrinogen, as well as reduced activation of pro-UPA. It also showed resistance to all three inhibitors. The 99-loop of FSAP is mainly composed of acidic residues, which are not conserved across different species. The single amino acid mutant E96S also exhibits reduced cleavage of histones, fibrinogen, and S2288 and is resistant to aprotinin but not TFPI and C1Inh. Residue 99 of this loop plays a crucial role in determining selectivity of the S2 pocket, and this residue is Tyr in FIXa, FXa, and tPA, Leu in thrombin, and Thr in APC(34). Interestingly, this residue is Glu in FSAP, which is unique in comparison to other serine proteases. The mutant D173A/E99S exhibited reduced catalytic activity against S2288, diminished cleavage of histones and fibrinogen, but only showed resistance to C1Inh. Our results clearly show that the 99-loop of FSAP has a significant role in its enzymatic specificity and functionality.

### Mutations in the Oxyanion hole stabilizing (186) loop

This region encompasses both the S1 pocket (Asp 189) and the oxyanion hole (Ser 195 and Gly 193), which is involved in catalysis. It has previously been demonstrated that trypsin with Q192D mutation is resistant to inhibition by aprotinin(35), and thrombin can be rendered sensitive to aprotinin by the E192Q mutation(36). We found that Q192D in SPD-FSAP is resistant to C1Inh, aprotinin, as well as TFPI, and is highly conserved across all species (Fig. S2). Lower histone cleavage was observed with this mutant, but the difference was not statistically significant. However, its activity towards S2288 and fibrinogen was preserved. Resistance to all inhibitors could increase the half-life of this mutant *in vivo*.

### Mutations in the S1 Entrance frame (220 loop)

Together with the 186 loop, the S1-entrance frame forms part of the S1 pocket and controls access of the substrate to the active site. It also plays a role in regulation by Na^+^ ions in FVIIa, thrombin, protein C, FIX, and FX (37), as well as FSAP(24). The MI-SNP (G221E) in FSAP prevents zymogen conversion, indicating the importance of the 220-loop for the activation of FSAP(38). In most proteases, position 218 is highly conserved with Gly, but in FSAP, this residue is Glu (Fig. S2). When mutated to Ala, it cleaved S2288, histones, and fibrinogen normally. However, it was partially resistant to inhibition by C1Inh but not by aprotinin or TFPI. These results indicate that E218 is specifically important for interaction with C1Inh. However, the K223S/R225A mutant in the same loop only exhibited resistance to aprotinin. These mutants can be used to investigate the regulation of FSAP activity by ions.

### Limitations of the study

The use of the recombinant SPD domain of FSAP for analyzing substrate and inhibitor specificity introduces a limitation that only short-range interactions within the SPD region are tested. FL-FSAP contains, in addition to the SPD, three EGF domains, a kringle domain, and an N-terminal region, and interactions of substrates and inhibitors with these domains likely play a major role in regulating its proteolytic activity. This is exemplified by its better inhibition by TFPI and C1Inh, but it cleaves fibrinogen and histone more poorly. Furthermore, direct binding studies with these mutants on the background of an activity-deficient enzyme are needed to distinguish between the effects of mutations on enzyme activity and direct binding. Finally, *in silico* interaction models between SPD-FSAP and its interacting partners can be used to understand the consequences of the mutations in a more precise manner. The results of these mutations have been largely discussed in the context of a direct interaction between the enzyme and its target. Additional interpretations, such as altered conformation of these mutations and protein stability, may also account for the observed effects. The use of aprotinin to titrate the active site concentration in proteins is not valid for mutants that exhibit altered interaction with the inhibitors, and alternative methods to measure this should have been used to consolidate the results.

## Conclusions

These findings contribute to the initial identification and characterization of specific regions within SPD-FSAP involved in interactions with its substrates and inhibitors. Further studies will help in creating mutants with altered properties to characterize the functions of FSAP.

## Acknowledgements

We thank Lukasz Wyrozemski for his technical assistance, Dr. Michael Etscheid, Paul Ehrlich Institute, Langen, Germany, for providing FL-FSAP purified from human plasma, and Prof. Tilman Hackeng, Maastricht University, Maastricht, Netherlands, for providing recombinant FL-TFPI.

## Author contributions

Sai Priya Sarma Kandanur: Investigation, Methodology, Formal analysis, Data Curation, Visualization, Writing - Review & Editing.

Bjørn Dalhus and Jonas Emsley: Supervision, Formal analysis, Writing – editing.

Sandip Kanse: Conceptualization, Supervision, Formal analysis, Writing - Original draft, Resources, Visualization.

## Data sharing statement

The data that support the findings of this study are available from the corresponding authors upon reasonable request.

---

**Fig. S1:**
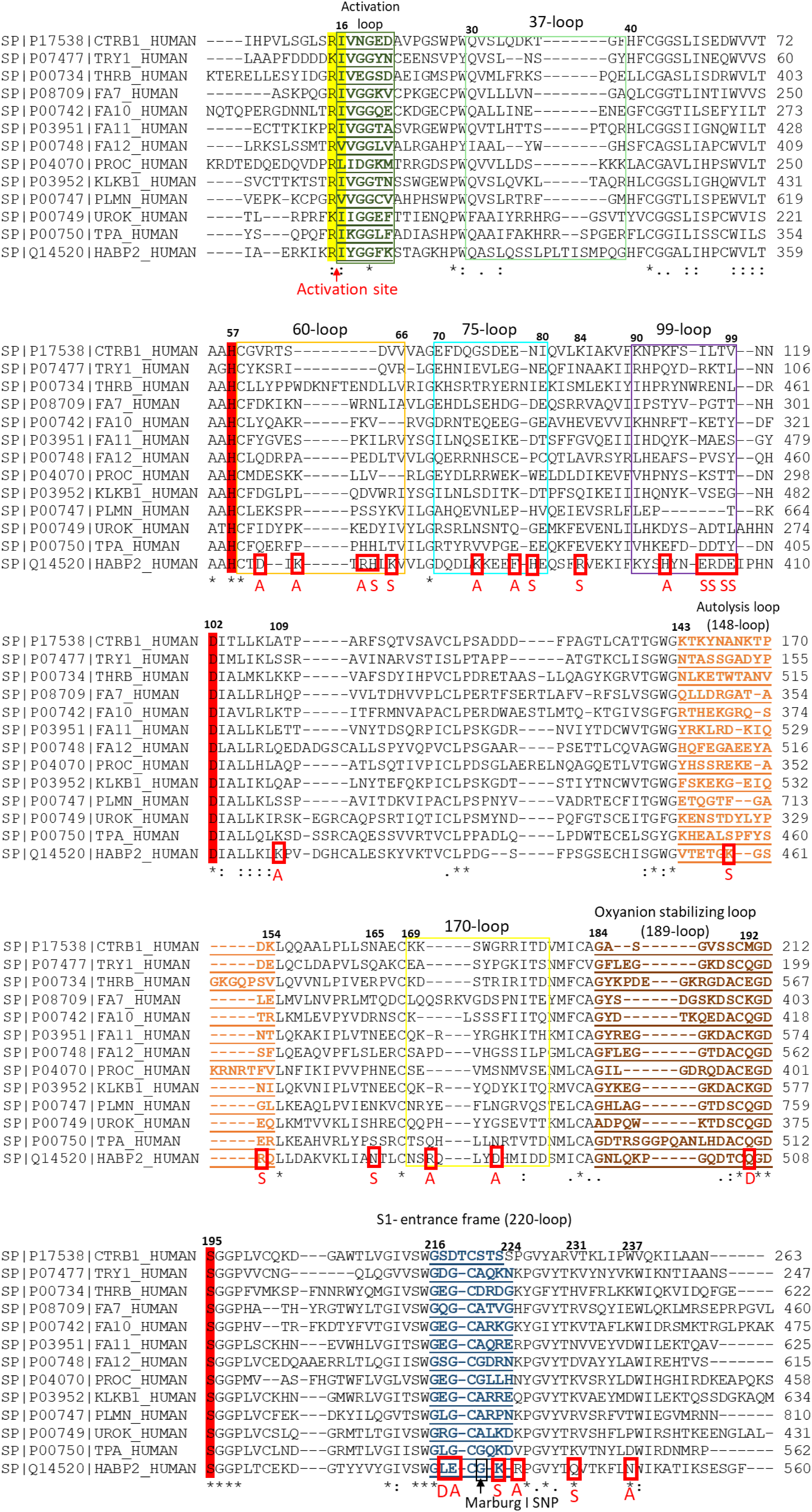
Sequence alignment of SPD-FSAP and related proteases: The serine protease domain of FSAP (SPD-FSAP) was aligned to its human paralogs, Chymotrypsin (CTRB1_HUMAN), Trypsin (TRY1_HUMAN), Thrombin (THRB_HUMAN), Factor VII (FA7_HUMAN), Factor X (FA10_HUMAN), Factor XI (FA11_HUMAN), Factor XII (FA12_HUMAN), A protein C (PROC_HUMAN), Kallikrien (KLKB1_HUMAN), Plasminogen (PLMN_HUMAN), urokinase-type plasminogen activator (UROK_HUMAN), and tissue-type plasminogen activator (TPA_HUMAN) using UniProt under default settings. The numbers in bold are based on the chymotrypsin numbering system. Numbers on the right represent amino acid position based on the individual protein sequence numbering. The catalytic triad His 57, Asp 102, and Ser 195 is highlighted in red, and the activation site is marked in yellow. The different surface loops are indicated, and mutations introduced in SPD-FSAP are marked with red boxes.

**Fig. S2:**
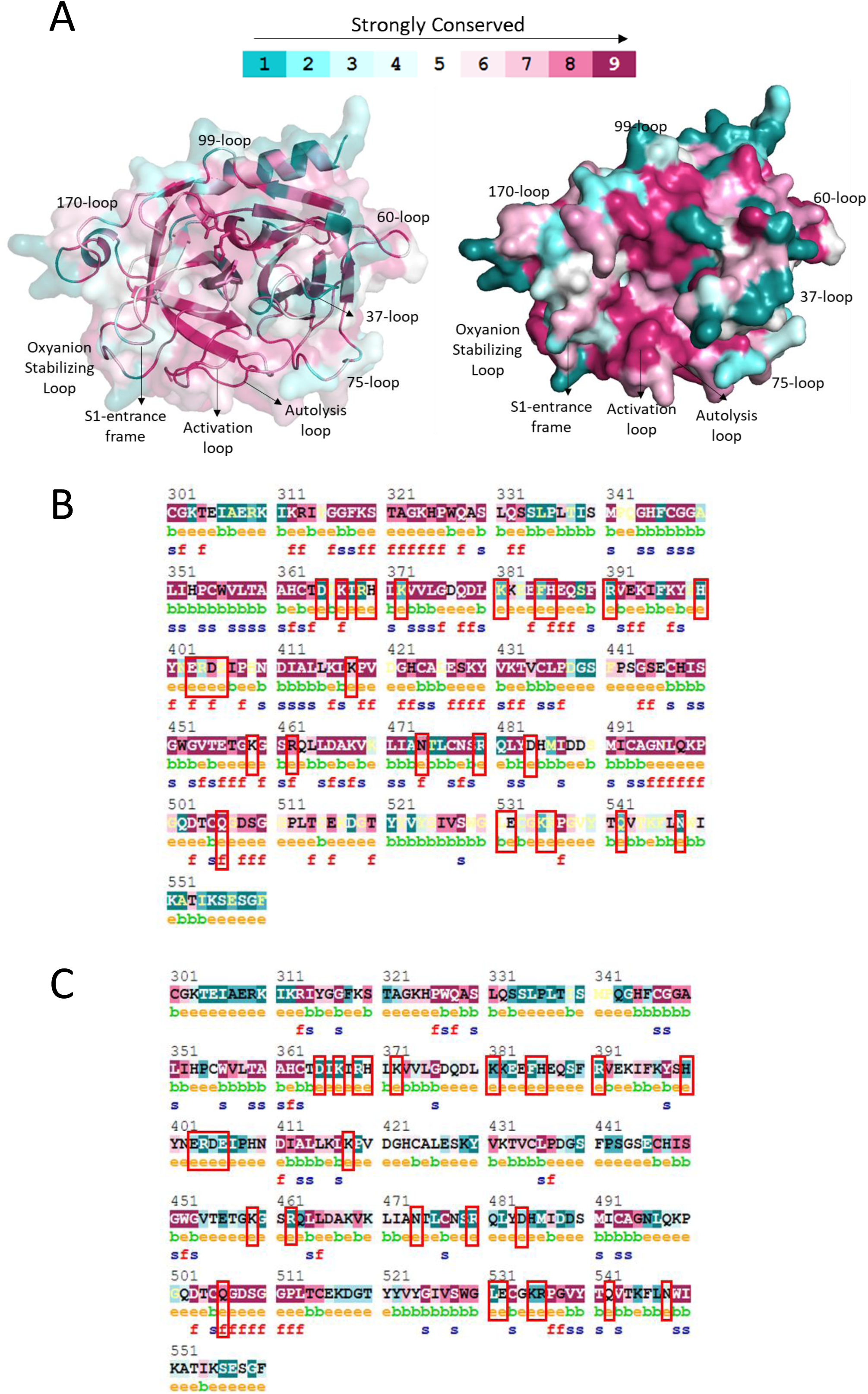
Conservation of SPD-FSAP (residues 313-560) across different species and human paralogs: **A.** Cartoon (left) and surface (right) representation of AlphaFold model of SPD-FSAP, created using ConSurf, depicting the conservation of amino acids across 81 different species of the subphylum vertebrates. **B.** The same data is shown as a sequence. **C.** Sequence conservation across 100 human paralogs. For both B and C, Residues that are exposed (e), buried (b), functional (f), structural (s), strongly conserved (purple), and variable (green) are predicted. Residues targeted for mutation are highlighted in red boxes. Numbering is based on the FSAP full-length sequence.

**Fig. S3:**
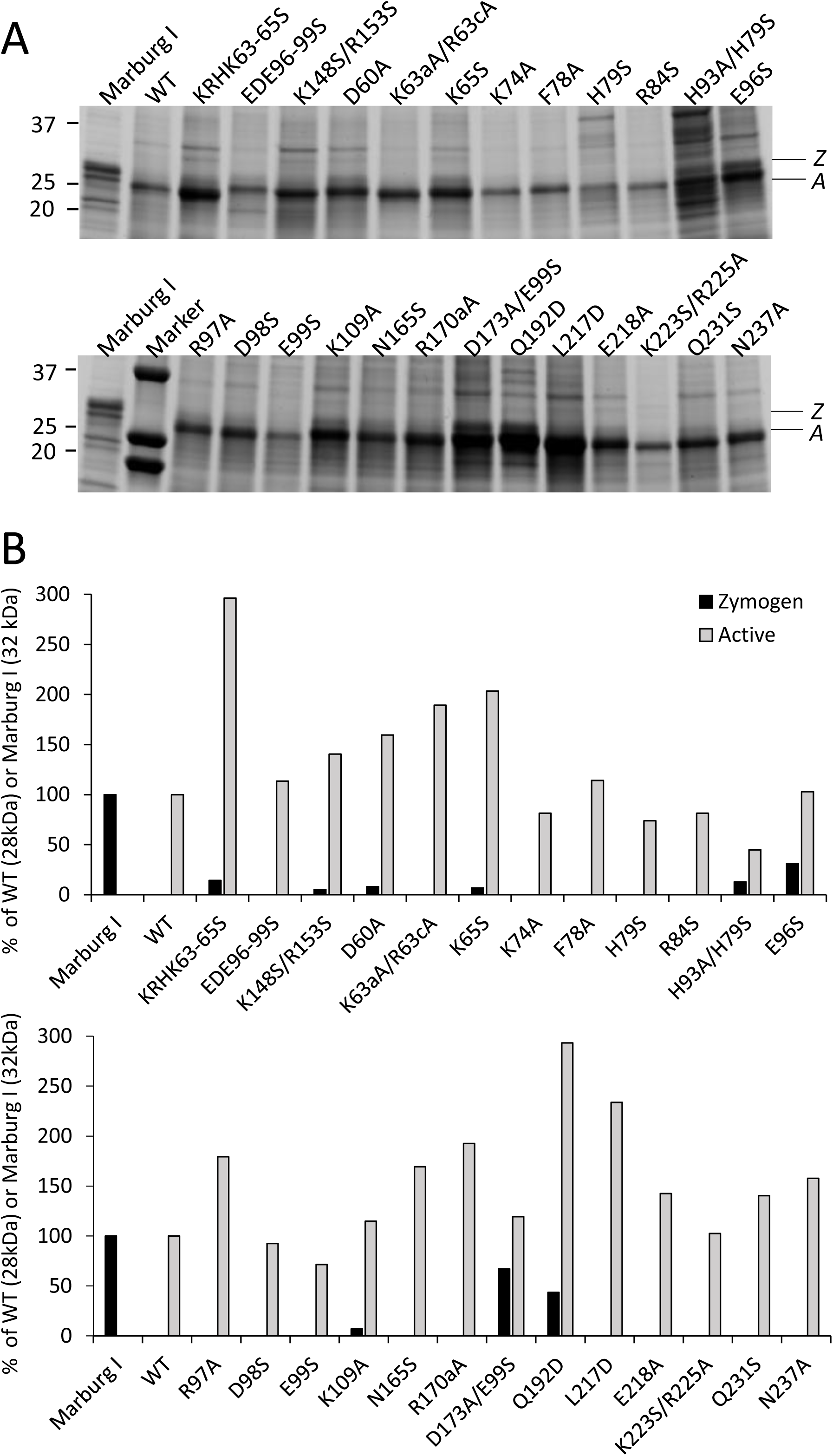
SDS-PAGE analysis of SPD mutants: **A.** Inclusion bodies were solubilized and refolded without purification on a Ni-NTA column. For each mutant, an equal volume of the sample was loaded on SDS-PAGE under reducing conditions. **B.** Densiometric analysis of the active (A) and zymogen (Z) fraction from each sample relative to the WT band (28 kDa) and the Marburg I band (32 kDa).

**Table S1:**
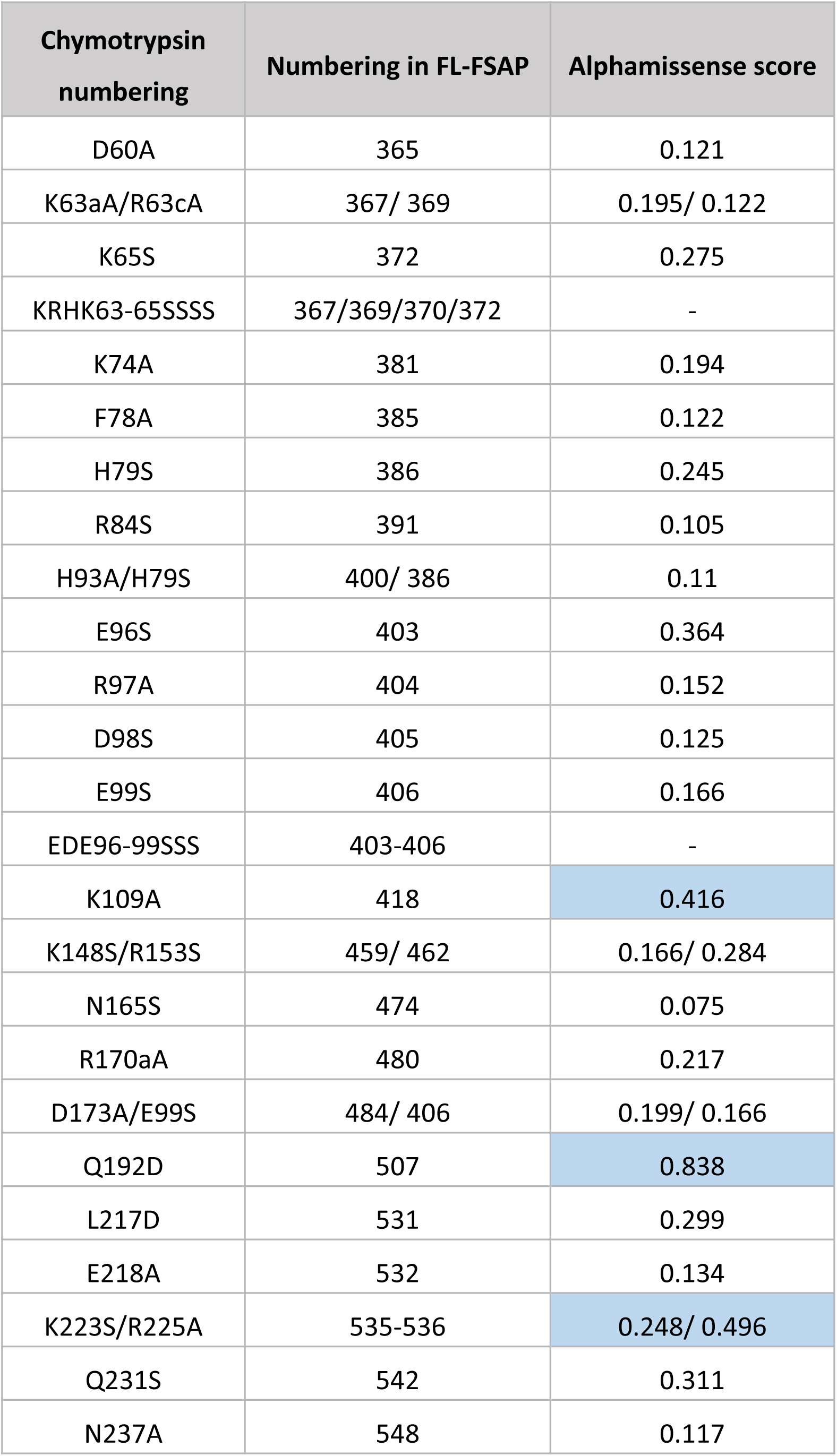
Chymotrypsin numbering and position of mutants in full-length (FL) FSAP (Uniprot Q14520). Alphamissence score from the Alphafold server for the likely pathogenicity of the mutation based on the evolutionary relationship between the amino acids at the different positions. Mutations that lead to a generally higher score are highlighted.

